# Is mechanical receptor ligand dissociation driven by unfolding or unbinding?

**DOI:** 10.1101/593335

**Authors:** Lukas F. Milles, Hermann E. Gaub

**Affiliations:** Lehrstuhl für Angewandte Physik and Center for Nanoscience, Ludwig-Maximilians-Universität, Amalienstr. 54, D-80799 München, Germany

## Abstract

Mechanical force can play a pivotal role in biological systems. Single Molecule Force Spectroscopy, is a powerful tool to probe the mechanics of proteins and their binding partners. Yet, it remains unclear how complex dissociation of a protein-protein interaction under mechanical forces occurs. Are receptor and ligand unbinding, or are they unfolding? We utilize an approach wherein receptor and ligand are expressed as a single molecule fused by a long flexible linker. Force is applied to the complex via an ultrastable handle. Consequently, the events during and following complex dissociation can be monitored. We investigate two high-affinity systems: The cohesin-dockerin type I interaction in which we find that a binding partner unfolds upon complex dissociation, and a colicin-immunity protein complex in which both proteins unfold completely upon unbinding. Mechanical receptor ligand dissociation thus can encompass unfolding of one or both binding partners.

## INTRODUCTION

Mechanical force can play a pivotal role in biological systems. Protein receptor-ligand complexes under mechanical stress have shown considerable resilience^1^, malleability^2^, and weakness^3^ against externally applied forces. In is various adaptations atomic force microscopy-based (AFM) single-molecule force spectroscopy (SMFS) has emerged as a powerful tool to uncover these behaviors^4–7^. Yet, it remains debatable in what pathway complex dissociation of a protein-protein interaction under mechanical forces occurs: Are these interactions unbinding, or unfolding? In conventional assays receptor and ligand are immobilized on different substrates, such as AFM cantilever and a surface. They are brought in contact until bound and subsequently dissociated by stretching until their complex breaks. Inherently, this method loses observation of binding partners *post* dissociation, as contact between cantilever and surface is lost when the connection severs. Thus, one lacks access to a more complex underlying mechanism reflected in e.g. intermediate states of unbinding, and what happens to the binding partners once dissociated. One possibility is that receptor and ligand unbind, *i.e.* they sever and remain two intact units. However, another conceivable pathway is the partial or complete unfolding of one or both of the binding partners. Especially high forces exceeding 50 pN routinely reach in AFM-based SMFS imply that protein domain unfolding is possible. As observation of binding partners is lost after dissociation in a conventional AFM-SMFS measurement, both pathways – unfolding and unbinding - will appear identical.

We use a single-molecule approach to experimentally address this issue, employing a system best described as a *tethered complex*. This polyprotein design strategy - not only suitable for AFM-SMFS as used in this study - consists of a receptor-ligand complex expressed as a single fusion protein. Successful implementations and descriptions of variations of this strategy have been established previously using DNA, peptide, or synthetic polymer linkers to connect receptor and ligand into a single unit or construct.^8–12^

Here, binding partners of the protein-protein complex of interest are expressed connected by a long, flexible, unstructured protein linker, around 30 nm in contour length. A reliable, ultrastable handle is used to apply force to the complex under investigation. This polyprotein assembly strategy is not only suitable for AFM-SMFS as demonstrated here, consists of a receptor-ligand complex expressed as a single fusion protein. Binding partners of the complex of interest are expressed connected by a long, flexible, unstructured protein linker (sequences in methods). A reliable, ultrastable handle is used to apply force to the complex under investigation^13^. As referenced above this was not the first installment of this concept.

Notably, the “ReaLism” implementation by Kim et al. established this strategy successfully for a protein-protein interaction: The “flex-bond” of the A1 domain of von Willebrand factor to the glycoprotein Ib α subunit in optical traps at forces below 30 pN.^8^ In a very similar approach in AFM-SMFS, the interaction between two Ig-like folds that connect the terminus of titin to obscurin 1 was investigated.^9^ A myomesin homodimer interaction has been probed by disulfide linkage of the two interacting domains through a mechanically compliant motif, again in AFM-SMFS, although by non-specifically tethering the protein.^11^

Building upon these advances regarding AFM-SMFS, here, these are extended using a specific handle that can withstand forces almost an order of magnitude larger to those mentioned above (up to 700 pN). Furthermore, longer linkers sequences are introduced and provided in a gene template for generalized construction of tethered complex systems.

As a benchmark complex of interest, we investigate the well-established cohesin-dockerin (coh-doc) type I interaction and reproduce the rupture forces exceeding 100 pN. Through the tethered complex approach we find that the dockerin domain unfolds upon unbinding, which offers an explanation for previous observations regarding calcium-dependent refolding.^14,15^ Moreover we probe the mechanical resilience of the extremely high-affinity (K_D_ ∼ fM) interaction of a bacteriocin (colicin E9) and its immunity protein (Im9). The tethered complex yields low receptor-ligand dissociation forces of around 60 pN. We can conclusively demonstrate that in a strict sense no unbinding takes places, as complex dissociation is rather a simultaneous unfolding of both receptor and ligand. Significantly, the tethered complex definitively settles the question of unfolding or unbinding in complex dissociation for these two model systems and confirms the concept of mechanical dissociation through domain unfolding.

## RESULTS AND DISCUSSION

The tethered complex principle rests upon the ultrastable type III CohesinE : Xmodule-dockerin (CohE:Xdoc) interaction from *Ruminococcus flavefaciens.* This complex exhibits a high affinity (K_D_ ∼ 20 nM) and a mechanical stability routinely reaching > 600 pN, exceeding the mechanical strength of almost all known complexes characterized in this regard.^6,13,16^ A second crucial step was to establish a very long, flexible protein linker sequence of tunable length to connect receptor and ligand in a single fusion protein. A soluble and sufficiently long, non-repetitive, easily clonable linker sequence is a crucial component of this system. Here, a flexible glycine-serine linker was combined with GSAT and SEG sequences (for sequences see methods). A molecular fingerprint is included in the system, to validate single-molecule and specific tethering via the CohE:Xdoc ultrastable handle.

Combining these components, we constructed a gene expressing a single polyprotein fusion comprised of a fingerprint domain of known unfolding force and pattern (Fig. 1) - here the ddFLN4 fold^17^. This small protein of around 100 amino acids unfolds at forces below 100 pN with a distinctive substep^18,19^, making it straightforward to identify in the force-extension data. ddFLN4 is followed by the receptor and ligand connected *via* a flexible polypeptide linker of adjustable length. The first binding partner will be tethered form its N-terminus, whereas the other binding partner will be force loaded form its C-terminus. The final, C-terminal domain is the Xmodule-dockerin handle. The Xdoc binding CohE construct on the cantilever contained a refolding Carbohydrate binding module (CBM) fingerprint^20^. All proteins were covalently bound immobilized on the surface via the ybbr-tag^21^. Through the ultrastable CohE:Xdoc tether one can apply force to the complex of interest, observing both its dissociation and all pathways succeeding, as the system is still tethered through the flexible linker. Only as the final event, following the unfolding of all protein domains of the complex of interest and the fingerprint, the dissociation of the ultrastable handle can be observed (See Fig. 2A).

**Figure 1.**
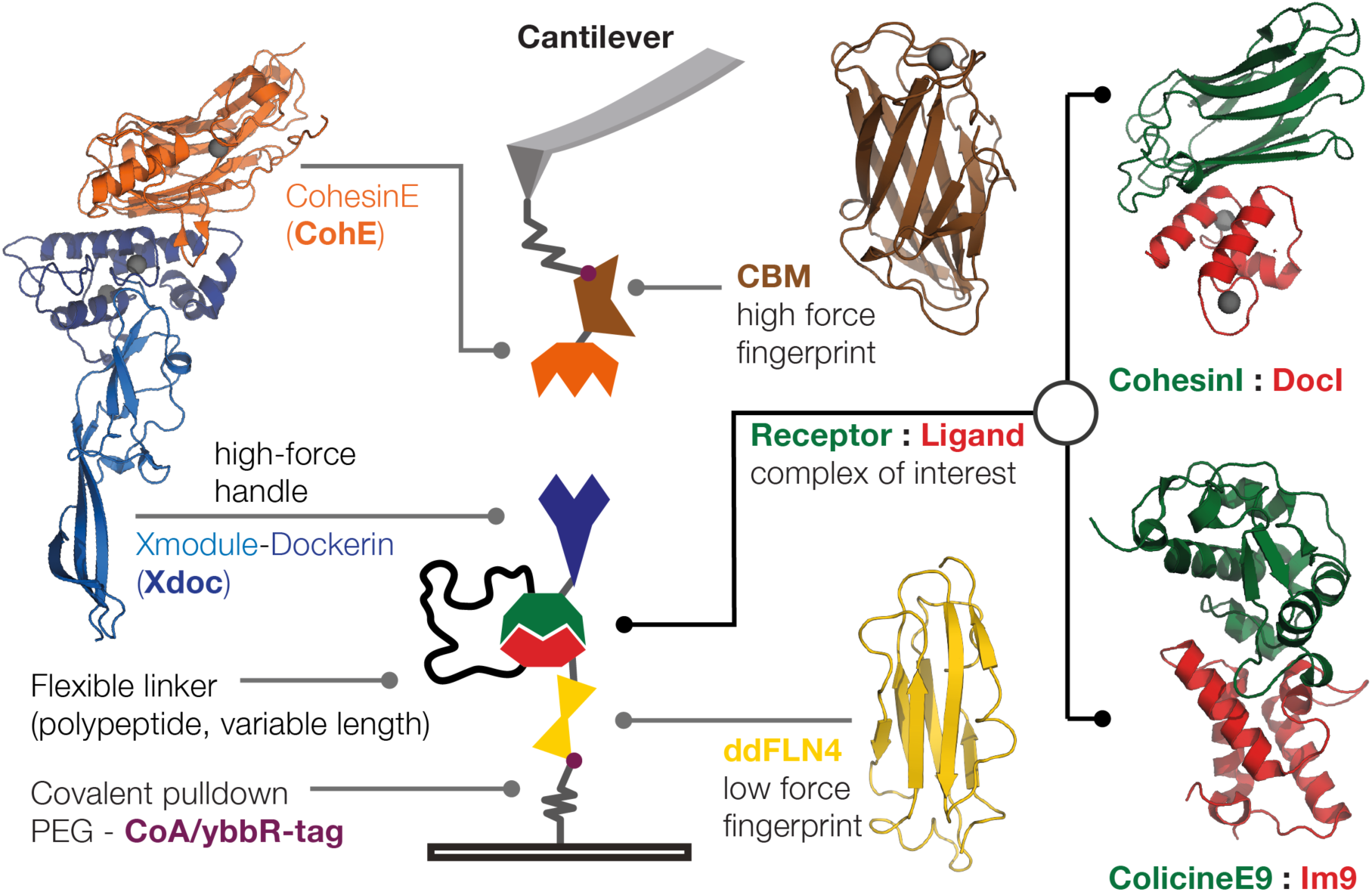
Overview over the tethered complex system components. Principle of a tethered complex, through fusion with known protein unfolding patterns the behavior of an unknown system may be probed. Receptor and ligand of interest (red/green) are connected by a flexible linker of programmable length. Here model complexes Coh-Doc type I (PDB 1OHZ) and colicinE9-Im9 (PDB 1EMV) were chosen. The construct is attached to the surface by a ybbr-tag and contains a refolding ddFLN4 fingerprint (yellow, PDB 1KSR). Force is applied to the system via an Xmodule-dockerin (Xdoc, blue, PDB 4IU3) which binds to its CohE binding partner (orange) on the cantilever, where a CBM (PDB 1NBC) fingerprint (brown) is included.

**Figure 2.**
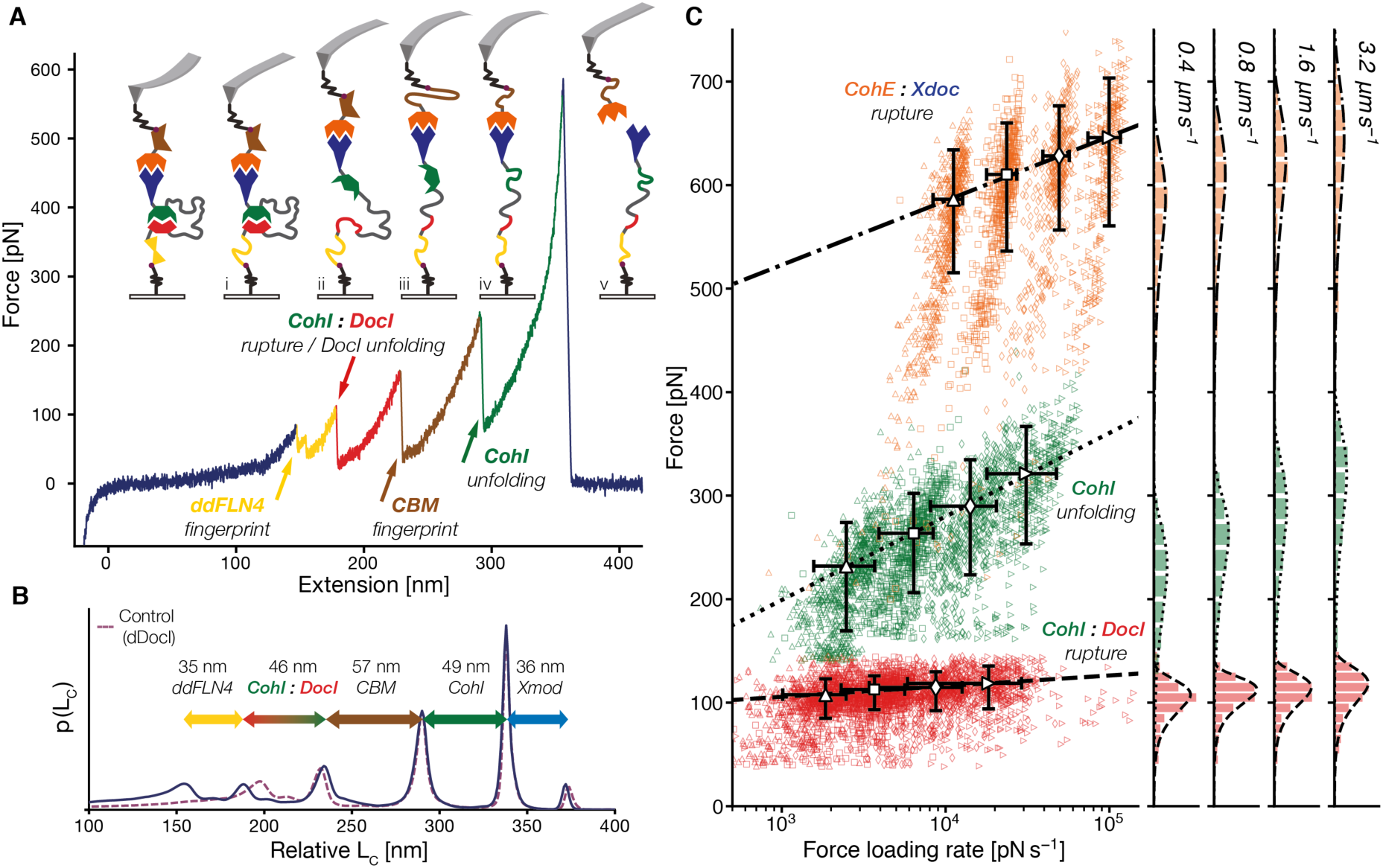
Tethered complex for cohesin-dockerin type I. (A) Force extension curve and corresponding representations of the force response of the system when probed by the ultrastable CohE-Xdoc type III interaction. After the unfolding of the low force fingerprint (subpanel (i), ddFLN4, with characteristic substep, in yellow) the CohI- DocI complex breaks with DocI unfolding (ii), ensued by the unfolding of the CBM fingerprint (brown) around 150 pN (iii). Consequently, the CohI domain (green) of the now severed Coh-Doc type I interaction unfolds here at around 225 pN (iv), and finally – after all components of the system are mechanically denatured, the ultrastable CohE:Xdoc interaction (orange:blue – respectively) breaks at ∼ 600 pN (v). (B) The aligned contour length diagram shows (blue line) that first the ddFLN4 fingerprint with its intermediate substep unfolds, yielding the expected around 35 nm. It is followed by the CohI-DocI interaction, where the contour length increment freed corresponds to the expected length of both the flexible linker, as well as the contour length of the unfolded DocI. Next, the CBM fingerprint with its characteristic 57 nm contour length increment unfolds, followed by the cohesin domain, which can be identified through its slightly shorter contour length of 49 nm. The dashed purple line shows the diagram for the dDocI control construct where the DocI was mutated out. Here, the 46 nm increment of receptor-ligand dissociation of Coh-Doc type I is clearly absent - as no CohI:DocI complex can form. The CBM, ddFLN4, and CohI unfolding contour lengths remain unchanged – as is to be expected. (C) The dynamic force spectrum (from retraction velocities of 0.4 to 3.2 µm s^-1^) for the ultrastable Xdoc-CohE handle (without Xmodule unfolded, orange, Δx = 0.15 nm, k_off_^0^ = 1.4E-7 s^-1^, N = 2043), the unfolding of the CohesinI domain (green, Δx = 0.12 nm, k_off_^0^ = 0.097 s^-1^, N = 2414), and the simultaneous unfolding and dissociation of the CohI:DocI complex of interest at around 100 pN (red, Δx = 0.93 nm, k_off_^0^ = 1.2E-8 s^-1^, N = 3363).

As a complex of interest, the well-established cohesin:dockerin type I (CohI:DocI) interaction was investigated^22,23^. Coh:Doc complexes assemble the cellulosome, an extracellular cellulose degrading enzyme network^24^. Previously observed rupture forces around 120 pN for CohI:DocI^25^ could be reproduced in the tethered complex system. Moreover, in the force extension curve and corresponding contour length diagrams^26^, both fingerprints, CBM and ddLFN4 could be identified (see Fig. 2B), confirming specific and single-molecule tethers. The remaining contour length increments were clear single events of 46 nm and 49 nm. The first 46 nm increment was associated with rupture forces around 100 pN, as is to be expected for CohI:DocI. The increment, however, matched the unfolding of the complete dockerin domain *plus* the expected freed contour length increment of the linker, not just the contour length of the flexible linker. The 49 nm unfolding event corresponds to the expected contour length increment for CohI unfolding and only occurs as the last step before CohE:Xdoc handle dissociation. These data strongly suggest that the dockerin domain unfolds upon unbinding from its cohesin partner. In a control construct where the DocI was deleted from the construct the 46 nm increment of DocI unbinding and unfolding was missing (Fig. 2B), strengthening the assignment of this increment to DocI:CohI dissociation. Therefore, DocI unfolds when mechanically dissociated from its CohI receptor.

Furthermore, these results offer an explanation for previous observations regarding calcium-dependent refolding of DocI. Dockerins have two important calcium binding loops, essential for their structural integrity^27^. Previously, it had been found that DocI immobilized on the AFM cantilever failed to interact with surface bound cohesin in the presence of calcium-chelating EDTA. However, the interaction could be recovered, when calcium was reintroduced.^25^ In light of the results presented here, clearly the dockerin unfolds upon unbinding, and could notrefold in the presence of EDTA. The cohesin domain unfolded later at forces ranging from 250 to 300 pN, implying that it remained intact after dissociation. These unfolding force determined here are also comparable to previous results.^28,29^

Moreover the mechanical resilience of the extremely high-affinity K_D_ ∼ fM interaction of a bacteriocin (colicin E9) and its immunity protein (Im9) was probed^30–32^. The E9:Im9 system is particularly suitable for the tethered complex approach: ColicineE9 is a fast endonuclease that kills bacteria rapidly by shredding their chromosome, recombinant expression in *E. coli* thus is challenging. The immunity protein Im9 binds to E9 with an extremely high affinity, effectively blocking its nuclease activity. By combining these to proteins in a tethered complex, E9 is permanently blocked by Im9, which is translated first by the ribosome in the construct. This “autoinhibited” system can be expressed recombinantly with good yield. The fingerprint domains unfold with their expected increments and forces (Fig 3 B,C). A single contour length increment of around 105 nm remains, which can – by exclusion - only be the E9:Im9 interaction. The tethered complex yields low receptor-ligand dissociation forces of around 60 pN (see Fig. 3), which is in good agreement with a previous study.^33^ The 105 nm increment matches E9 unfolding *plus* Im9 unfolding *plus* linker contour length. Thus, here, both binding partners completely unfold upon dissociation - noticeable in a clean single stretch contour length increment, which has not directly observable substructure – at least within the AFM’s force resolution. In a strict sense, thus, this system is not mechanically dissociated but completely unfolded, whereby the interaction is separated.

**Figure 3.**
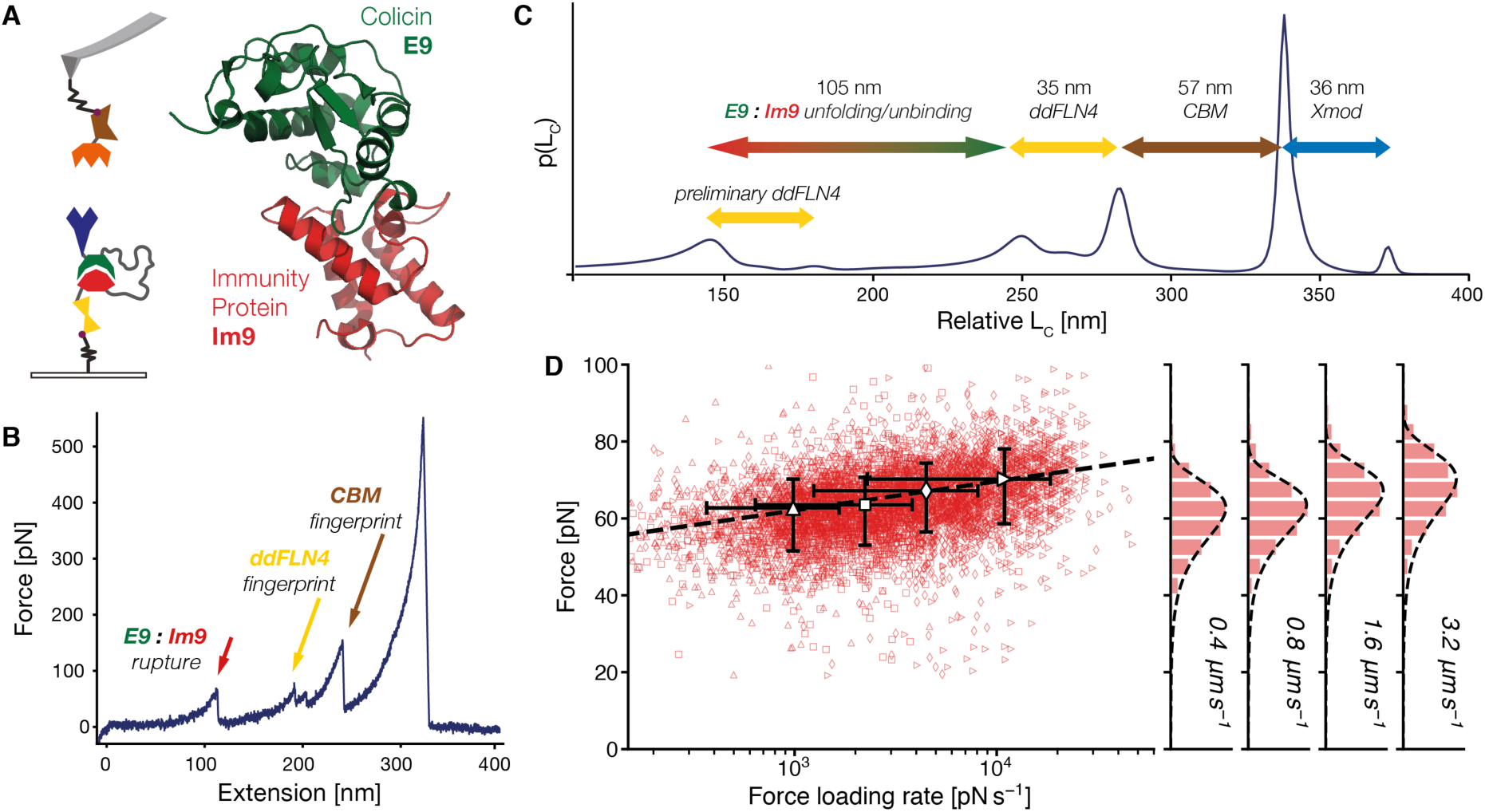
Tethered complex for colicin E9 and its immunity protein Im9. (A) Setup with structure of Im9:E9 (red:green) in the tethered complex system. (B) An exemplary force extension trace for the system. In contrast to Fig. 2, here the weaker Im9:E9 interaction breaks at already 60 pN (red arrow), weaker than the unfolding forces of both fingerprint (ddFLN4 and CBM, yellow and brown arrows respectively), the last event is the dissociation of Xdoc:CohE. (C) A corresponding contour length diagram at the bottom demonstrates the unfolding pathway. The weak E9:Im9 interaction usually breaks first unfolding both binding partners in the process, as the contour length increment produced by their unbinding adds up to both their domain lengths and the length of the linker region (∼ 105 nm in total). Next, the ddFLN4 fingerprint unfolds in its characteristic two step unfolding pathway. The order of these two may be flipped as their rupture forces are very similar, which is noted in the contour length diagram as some curve show the ddFLN4 unfolding before Im9:E9 rupture (preliminary ddFLN4). (D) The resulting dynamic force spectrum (from retraction velocities of 0.4 to 3.2 µm s^-1^) for the E9:Im9 complex (Δx = 1.3 nm, k_off_^0^ = 2.0E-6 s^-1^, N = 5879) unfolding and unbinding with a noticeably very flat log-linear slope of the most probable rupture force scaling with the force loading rate.

As elaborated in the introduction this was not the first implementation of this strategy. Here the focus was on the unfolding of binding partners of a receptor-ligand complex during its dissociation. However, there are other applications and advantages to using a tethered complex: Usually, only a single-receptor ligand system can be probed with a single cantilever. Only a one protein species can be fused reliably to the tip. Using tethered complexes, one can probe many different interactions with the same high-strength handle on the cantilever by swapping the tethered complex on the surface^29^. In addition, if the ultrastable handle is very reliable - as the CohE:Xdoc - it can serve as a proxy for non-refolding and sensitive complexes. These can be tested in a tethered complex, ensuring that a fresh, completely unprobed interaction is dissociated every time. This is in contrast to conventional force spectroscopy where the biomolecules on the cantilever remain unchanged through the course of the experiment, a disadvantage if they are susceptible to cantilever wear or not refolding. In turn, this feature also enables an experimental approach to address the lack of comparability of different receptor-ligand systems with a single force probe, as calibration uncertainties between AFM cantilevers make comparisons between their data challenging^34^. Usually, only a single-receptor ligand system can be probed with a single cantilever, as only a single protein species can be fused to the tip in a controlled manner, if not orthogonal fingerprinting is used^35^. Using tethered complexes, one can probe many different interactions with the same high-strength handle on the cantilever. Thus, different receptor-ligand complexes can be compared in absolute force with a single force probe. Furthermore, if a single complex is tethered it could be probed many times by keeping the high-force tether and reapproaching the surface to reform the complex and dissociate it multiple times, acquiring a dynamic force spectrum from one single molecule.^36^

Both systems investigated here have extremely high binding affinities (K_D_: CohI:DocI around pM, Im9:E9 around fM) so they will always be in the bound state when probed by force, this is reflected in the force traces which almost exclusively show complex rupture events. For lower affinity systems one could distinguish between bound, and unbound states when the dissociation contour length increment is missing. While the linker sets up receptor and ligand at a very high local concentration relative to each other, a rough estimate of complex fraction bound should be possible with this approach. Also, the close proximity induced by the linker will allow for the probing of very slow on-rate complexes that could not be probed by conventional AFM-SMFS. Especially for homodimers with high affinities, that could not be separated onto cantilever and surface without still being in a dimer state, the tethered complex approach provides a straightforward means to probe their mechanics. Another advantage of this strategy consists of the simultaneous determination of unfolding forces of the individual components of the system, when determining their complex rupture force. Here the unfolding force of the CohI from *C. thermocellum* was determined to be around 250 pN - merely as a byproduct of the probing strategy. The tethered complex approach may now be generalized with even higher stability handles such as the hyperstable SdrG:Fgβ interaction and its homologs, which merely requires a short (< 15 amino acid) peptide target as a hyperstable handle – greatly reducing tethered complex construct size and complexity.^37–39^

## CONCLUSION

We conclusively demonstrate that in a strict sense no unbinding takes places, when E9:Im9 and Coh-Doc type I are separated mechanically. Complex dissociation is rather a simultaneous unfolding of one or both domains of receptor and ligand. The tethered complex approach contributes to the debate of unfolding or unbinding in mechanical complex dissociation for these model systems, establishing that unbinding and unfolding can be connected. More interactions may be tested and compared with this design strategy and the linker sequence validated here. Hints at the biological functions of unfolding upon mechanical dissociation may provide more detailed perspectives on the fundamental mechanobiology and molecular mechanisms of protein-protein interactions.

## ACKNOWLEDGMENTS

We would like to thank Wolfgang Ott, Markus A. Jobst, Ellis Durner, and Tobias Verdorfer for helpful discussions and instrument development. We gratefully acknowledge funding from an advanced grant of the European Research Council (Cellufuel Grant 294438), and from the Deutsche Forschungsgemeinschaft SFB 863 and SFB 1032.

## METHODS

### Gene construction and protein expression

The carbohydrate binding module (CBM) and cohesin type I (CohI) genes are part of CipA, from *C. thermocellum*. The cohesin type III (CohE) from ScaE and Xmodule-dockerin (XDoc) from Ctta are genes of *R. flavefaciens FD-1*. All of the above were kind gifts of E. A. Bayer (Weizmann Institute, Rehovot, Israel). The DocI gene was amplified from the C. thermocellum genome (ATCC 27405, from DSMZ 1237). The *Dictyostelium discoideum* 4^th^ filamin domain (ddFLN4), the Colicine E9 (E9) and immunity protein 9 (im9) as well as the long linker region genes were synthesized codon-optimized for expression in *Escherichia Coli* as a linear DNA fragment (GeneArt – ThermoFisher Scientific, Regensburg, Germany). The linker domain was sourced partially from the iGEM parts databank (http://parts.igem.org : BBa_K404300, BBa_K243029 both from Freiburg Bioware Teams 2009/2010). The full protein linker sequence is: 73 amino acid linker (contour length ∼ 27 nm) for the Im9/E9 construct: GGGSAGGSGSGSSGGSSGASGTGTAGGTGSGSGTGSGGGSGGGSEGGGSEGGGSEGGGSEGGGSEGGGSGGGS Another combined linker assembled from GS and native ScaA cellulosome linkers reads (63 amino acids, ∼ 23 nm contour length) for the DocI/CohI construct: DKFPVAENPSSHPWTSASGSGSGTAEGGSTAGSVVPSTQPVTTPPATTKPPATTIPPSDDPNA And an extended version of ∼ 44 nm contour length: DKFPVAENPSSGGGSAGGSGSGSSGGSSGASGTGTAGGTGSGSGTGSGGGSGGGSEGGGSEGGGSEGGGSEGGGSEGGGSGGGSESTAGSVVPSTQPVTTPPATTKPPATTIPPSDDPNA All plasmids were assembled using the Gibson assembly strategy^40^ (New England Biolabs, MA, USA) into pET28a Vectors. The C63S mutation in the CBM and C18S mutation in the ddFLN4 had been introduced previously by blunt end ligation cloning using T4 Ligase (Thermo Scientific, MA, USA). Final open reading frames of all constructs were checked by DNA sequencing (Eurofins Genomics, Ebersberg, Germany).

### Protein expression and purification

All proteins were expressed ybbr-tagged^21^ in *E. Coli* NiCo21(DE3) (New England Biolabs, MA, USA). Precultures of 5 mL in LB medium, grown overnight at 37° C, were inoculated in ZYM-5052 autoinduction media containing 50 μg/mL kanamycin and grown for 6 h at 37° C and then 24 h at 24° C.^41^ Bacteria were spun down into a pellet, to be stored frozen at −80° C. The pellet was resuspended in Lysis Buffer 100 μg/mL Lysoszyme and 10 μg/mL DNAseI (Roche) were added, and cells were lysed through sonication (Bandelin Sonoplus GM 70, Tip: Bandelin Sonoplus MS 73, Berlin, Germany at 50% power, 30% cycle 2 × 10 min) followed by centrifugation at 38000 g for 45 minutes. The supernatant was applied to a Ni-NTA column (GE Healthcare, MA, USA) for HIS-Tag purification and washed (25 mM TRIS, 500 mM NaCl, 20 mM Imidazole, 0.25 % (v/v) Tween20, 10 % (v/v) Glycerol), extensively. The protein was eluted with 200 mM imidazole. Protein containing fractions were concentrated over regenerated cellulose filters (Amicon, Merck, Darmstadt, Germany), exchanged into measurement buffer (TBS-Ca: 25 mM Tris, 72 mM NaCl, 1mM CaCl_2_) by polyacrylamide columns (Zeba, Thermo Scientific, MA, USA), and frozen with 25 % (v/v) glycerol in liquid nitrogen to be stored at −80° C until used in experiments. Protein concentrations were measured with spectrophotometry with typical yields of more than 100 μM (on a NanoDrop 1000, Thermo Scientific, DE, USA).

### AFM sample preparation

An exhaustive AFM-SMFS protocol has been published previously.^42^ AFM Cantilevers (Biolever Mini, Olympus, Tokyo, Japan) and cover glass surfaces are modified with aminosilane identically. In brief, after UV-Ozone cleaning, surfaces were incubated in (3-aminopropyl)-dimethyl-ethoxysilane (APDMES, abcr, Karlsruhe, Germany) baked at 80° C for 1 h and stored overnight under argon. Both surfaces were covered with 5 kDa heterobifunctional Succinimide-PEG-Maleimide (Rapp Polymere, Tübingen, Germany) dissolved in 50 mM HEPES (pH 7.5) for 30 min. After rinsing with ultrapure water, 1 mM Coenzyme A in a 50 mM sodium phosphate pH 7.2, 50 mM NaCl, 10 mM EDTA buffer was applied for 1 h. The protein samples were exchanged into TBS-Ca supplemented with 10 mM MgCl_2_. After rinsing in water again, the cantilevers were incubated with 50 μM CohE-CBM-ybbr and 28 μM Sfp phosphopantetheinyl transferase (SFP) for 2 h. The glass surfaces were incubated with 0.5-2 μM tetherd complex constructs and 13 μM SFP for 30 min. Both samples were rinsed extensively with at least 60 mL TBS-Ca before measurement.

### AFM-SMFS

AFM-SMFS data was acquired on a custom-built AFM operated in closed loop by a MFP3D controller (Asylum Research, Santa Barbara, CA, USA) programmed in Igor Pro 6 (Wavemetrics, OR, USA). Cantilevers were briefly brought in contact with the functionalized surface and then retracted at constant velocities of 400, 800, 1600, and 3200 nm/s. Following each curve, the glass surface was moved horizontally by 100 nm to expose an unused surface area. Typically, 50000 - 100000 curves were recorded per experiments. When quantitative comparisons of forces were needed a single cantilever was used to probe multiple surfaces. Cantilevers were calibrated using the equipartition theorem method with typical spring constants between 50-110 pN nm^-1^.^43^

### SMFS data analysis

Data analysis was carried out in Python 2.7 (Python Software Foundation).^44–46^ Raw data were transformed from photodiode and piezo voltages into physical units with the cantilever calibration and piezo sensitivity. Laser spot drift on the cantilever relative to the calibration curve was corrected *via* the baseline noise for all curves. The last rupture peak was detected and the subsequent 20 nm were used to set the force baseline to zero. The origin of extension was then set as the first and closest point to zero force. A correction for cantilever bending given through the forces measured was applied to the extension datapoints. For peak detection, data were denoised with Total Variation Denoising (TVD, denoised data not shown)^47,48^, and rupture events detected as significant drops in force. A three regime model by Livadaru et. al.^49^ was used to model the elastic behavior of free contour lengths and transform into contour length space^26^ (with: stiff element b = 0.11 nm and bond angle γ = 41°). Peaks were assigned their contour length in diagrams assembled through Kernel Density Estimates (KDE) of the contour length transformed force-extension data. The KDE bandwidth was chosen as 1 nm. The loading rate was fitted as the linear slope of the last 4 nm preceding a peak.

Rupture force histograms for the respective peaks and dynamic force spectra were assembled from all curves showing the ddFLN4 and the CBM fingerprint. The most probable loading rate was determined with a KDE, with the bandwidth chosen by the Silverman estimator.^50^ This value was used to fit the unfolding or rupture force histograms with the Bell-Evans model for each pulling velocity.^51,52^ Errors are given as the asymmetric full width at half maximum (FWHM) of each probability distribution. A final fit was performed through the most probable rupture forces and loading rates for each pulling velocity to determine the distance to the transition state Δx_0_ and natural off-rate at zero force k_off,0_.

## SUPPLEMENTARY INFORMATION

### Full protein construct sequences for tethered complexes

ybbr-HIS-ddFLN4(C18S)-DocI(1ohz)-linker-CohI-Xdoc

*CohI:DocI construct with 63 amino acid linker with parts from ScaA*

MDSLEFIASKLAHHHHHHGSADPEKSYAEGPGLDGGESFQPSKFKIHAVDPDGVHRTDGGDGFVVTIEGPAPVDPVMVDNGDGTYDVEFEPKEAGDYVINLTLDGDNVNGFPKTVTVKPAPGSGSVIEGPAPQPTQPPVLLGDVNGDGTINSTDLTMLKRSVLRAITLTDDAKARADVDKNGSINSTDVLLLSRYLLRVIDKFPVAENPSSHPWTSASGSGSGTAEGGSTAGSVVPSTQPVTTPPATTKPPATTIPPSDDPNAGSDGVVVEIGKVTGSVGTTVEIPVYFRGVPSKGIANCDFVFRYDPNVLEIIGIDPGDIIVDPNPTKSFDTAIYPDRKIIVFLFAEDSGTGAYAITKDGVFAKIRATVKSSAPGYITFDEVGGFADNDLVEQKVSFIDGGVNVGNATVVPNTVTSAVKTQYVEIESVDGFYFNTEDKFDTAQIKKAVLHTVYNEGYTGDDGVAVVLREYESEPVDITAELTFGDATPANTYKAVENKFDYEIPVYYNNATLKDAEGNDATVTVYIGLKGDTDLNNIVDGRDATATLTYYAATSTDGKDATTVALSPSTLVGGNPESVYDDFSAFLSDVKVDAGKELTRFAKKAERLIDGRDASSILTFYTKSSVDQYKDMAANEPNKLWDIVTGDAEEE

ybbr-HIS-ddFLN4(C18S)-DocI(1ohz)-linker48nm-CohI-Xdoc

*same as above with longer linker*

MDSLEFIASKLAHHHHHHGSADPEKSYAEGPGLDGGECFQPSKFKIHAVDPDGVHRTDGGDGFVVTIEGPAPVDPVMVDNGDGTYDVEFEPKEAGDYVINLTLDGDNVNGFPKTVTVKPAPGSGSVIEGPAPQPTQPPVLLGDVNGDGTINSTDLTMLKRSVLRAITLTDDAKARADVDKNGSINSTDVLLLSRYLLRVIDKFPVAENPSSGGGSAGGSGSGSSGGSSGASGTGTAGGTGSGSGTGSGGGSGGGSEGGGSEGGGSEGGGSEGGGSEGGGSGGGSESTAGSVVPSTQPVTTPPATTKPPATTIPPSDDPNAGSDGVVVEIGKVTGSVGTTVEIPVYFRGVPSKGIANCDFVFRYDPNVLEIIGIDPGDIIVDPNPTKSFDTAIYPDRKIIVFLFAEDSGTGAYAITKDGVFAKIRATVKSSAPGYITFDEVGGFADNDLVEQKVSFIDGGVNVGNATVVPNTVTSAVKTQYVEIESVDGFYFNTEDKFDTAQIKKAVLHTVYNEGYTGDDGVAVVLREYESEPVDITAELTFGDATPANTYKAVENKFDYEIPVYYNNATLKDAEGNDATVTVYIGLKGDTDLNNIVDGRDATATLTYYAATSTDGKDATTVALSPSTLVGGNPESVYDDFSAFLSDVKVDAGKELTRFAKKAERLIDGRDASSILTFYTKSSVDQYKDMAANEPNKLWDIVTGDAEEE

ybbr-HIS-ddFLN4(C18S)-linker-CohI-Xdoc

*same as above control (no DocI domain)*

MDSLEFIASKLAHHHHHHGSADPEKSYAEGPGLDGGECFQPSKFKIHAVDPDGVHRTDGGDGFVVTIEGPAPVDPVMVDNGDGTYDVEFEPKEAGDYVINLTLDGDNVNGFPKTVTVKPAPGSGSVIEGPAPQPTQPPDKFPVAENPSSHPWTSASGSGSGTAEGGSTAGSVVPSTQPVTTPPATTKPPATTIPPSDDPNAGSDGVVVEIGKVTGSVGTTVEIPVYFRGVPSKGIANCDFVFRYDPNVLEIIGIDPGDIIVDPNPTKSFDTAIYPDRKIIVFLFAEDSGTGAYAITKDGVFAKIRATVKSSAPGYITFDEVGGFADNDLVEQKVSFIDGGVNVGNATVVPNTVTSAVKTQYVEIESVDGFYFNTEDKFDTAQIKKAVLHTVYNEGYTGDDGVAVVLREYESEPVDITAELTFGDATPANTYKAVENKFDYEIPVYYNNATLKDAEGNDATVTVYIGLKGDTDLNNIVDGRDATATLTYYAATSTDGKDATTVALSPSTLVGGNPESVYDDFSAFLSDVKVDAGKELTRFAKKAERLIDGRDASSILTFYTKSSVDQYKDMAANEPNKLWDIVTGDAEEE

ybbr-HIS-ddFLN4-im9-longlink-colicine9-Xdoc

*Im9:E9 construct with 73 amino acid linker*

MDSLEFIASKLAHHHHHHGSADPEKSYAEGPGLDGGECFQPSKFKIHAVDPDGVHRTDGGDGFVVTIE GPAPVDPVMVDNGDGTYDVEFEPKEAGDYVINLTLDGDNVNGFPKTVTVKPAPGSGSVIEGPAPQPTQ PPELKHSISDYTEAEFLQLVTTICNADTSSEEELVKLVTHFEEMTEHPSGSDLIYYPKEGDDDSPSGI VNTVKQWRAANGKSGFKQGGGSAGGSGSGSSGGSSGASGTGTAGGTGSGSGTGSGGGSGGGSEGGGSE GGGSEGGGSEGGGSEGGGSGGGSESKRNKPGKATGKGKPVGDKWLDDAGKDSGAPIPDRIADKLRDKE FKSFDDFRKAVWEEVSKDPELSKNLNPSNKSSVSKGYSPFTPKNQQVGGRKVYELHHDKPISQGGEVY DMDNIRVTTPKRHIDIHVVPNTVTSAVKTQYVEIESVDGFYFNTEDKFDTAQIKKAVLHTVYNEGYTG DDGVAVVLREYESEPVDITAELTFGDATPANTYKAVENKFDYEIPVYYNNATLKDAEGNDATVTVYIG LKGDTDLNNIVDGRDATATLTYYAATSTDGKDATTVALSPSTLVGGNPESVYDDFSAFLSDVKVDAGK ELTRFAKKAERLIDGRDASSILTFYTKSSVDQYKDMAANEPNKLWDIVTGDAEEE

